# A label-free approach for relative spatial quantitation of c-di-GMP in microbial biofilms

**DOI:** 10.1101/2023.10.10.561783

**Authors:** Catherine S. McCaughey, Michael A. Trebino, Allyson McAtamney, Ruth Isenberg, Mark J. Mandel, Fitnat H. Yildiz, Laura M. Sanchez

## Abstract

Microbial biofilms represent an important lifestyle for bacteria and are dynamic three dimensional structures. Cyclic dimeric guanosine monophosphate (c-di-GMP) is a ubiquitous signaling molecule that is known to be tightly regulated with biofilm processes. While measurements of global levels of c-di-GMP have proven valuable towards understanding the genetic control of c-di-GMP production, there is a need for tools to observe the local changes of c-di-GMP production in biofilm processes. We have developed a label-free method for the direct detection of c-di-GMP in microbial colony biofilms using matrix-assisted laser desorption ionization mass spectrometry imaging (MALDI-MSI). We applied this method to the enteric pathogen *Vibrio cholerae*, the marine symbiont *V. fischeri*, and the opportunistic pathogen *Pseudomonas aeruginosa* PA14 and detected spatial and temporal changes in c-di-GMP signal that accompanied genetic alterations in factors that synthesize and degrade the compound. We further demonstrated how this method can be simultaneously applied to detect additional metabolites of interest in a single experiment.

## Introduction

Cyclic dimeric guanosine monophosphate (c-di-GMP) is a universally important second messenger that regulates exopolysaccharide production, virulence, antimicrobial tolerance, flagellar motility, cell morphology, and cell cycle control across nearly all bacterial phyla.^1–3^ C-di-GMP is synthesized by diguanylate cyclases (DGCs) and hydrolyzed by phosphodiesterases (PDEs). DGCs contain a GGDEF catalytic domain for cyclizing two molecules of guanosine triphosphate (GTP), while PDEs contain either an EAL or HD-GYP domain, which hydrolyze c-di-GMP to the linearized form (pGpG) or two molecules of guanosine monophosphate (GMP), respectively.^4^ The regulatory role of c-di-GMP is particularly important in biofilm-forming bacteria, as evidenced by the high number of DGC and PDE enzymes present in the genomes of biofilm-forming organisms such as *Pseudomonas aeruginosa, Vibrio cholerae,* and *Vibrio fischeri* including many enzymes with dual DGC/PDE activity.^1,5,6^

There are a variety of methods available to detect c-di-GMP in biological samples. LC-MS/MS analysis is currently the most accurate and sensitive method for detecting c-di-GMP, particularly in liquid extracts of nutrient broths, which provide a cumulative amount of c-di-GMP from diverse populations of bacterial cells. Fluorescent detection of c-di-GMP is also commonly used on bulk liquid samples and solid samples, and has many advantages in efficiency of detection and application engineered *in vivo* systems.^5^ Numerous innovative approaches for the detection of c-di-GMP using fluorescent biosensors have been recently developed including fluorescent reporter systems,^7,8^ riboswitch-based biosensors,^9,10^ FRET-based biosensors,^11^ and others.^12–14^ In general, fluorescent detection methods are applied to bulk samples but can also be used to detect the spatial and temporal dynamics of c-di-GMP signaling in biofilms.^7,15^ However, these fluorescence methods rely on engineered strains or reporters that do not allow for a facile extension to environmental isolates or emerging pathogens and are indirect measures of c-di-GMP levels.

Generally, high c-di-GMP levels induce the expression of biofilm-related genes, resulting in an overproduction of extracellular matrix and a wrinkled phenotype in surface-grown colonies, however, there are also inconsistencies in this model.^4,16,17^ Particularly in organisms harboring a large number of DGC and PDE domains, which control different c-di-GMP-dependent processes and effectors, it is often seen that deletion or overexpression of single DGC or PDE genes results in unexpected changes to the global c-di-GMP levels.^16^ Biofilms have heterogeneous structures and chemical gradients that change dynamically over time and influence the regulation of these DGC and PDE enzymes.^18,19^ MS imaging (MSI) is an analytical technique that allows for the spatial detection of small molecules (100 – 2000 Da) in biological samples, including tissues, whole organisms, and bacterial samples grown on a variety of substrates.^20^ MSI can achieve an image resolution as low as 10-20 μm and enables the robust detection of thousands of chemical species in a single experiment.^21^ This label-free technique has a significant advantage over fluorescent detection techniques in that both known and unknown chemical species can be correlated spatially with biofilm formation and c-di-GMP metabolism. At present, microbial MSI requires optimization for different microbial cultures and analyte classes.^21^

Here we present an MSI technique for the direct detection of c-di-GMP in bacterial biofilm colonies grown on solid agar media using matrix assisted laser desorption/ionization (MALDI) mass spectrometry imaging (MSI). We validated our MSI technique using a fluorescent riboswitch in *V. cholerae* and we further applied the MSI detection of c-di-GMP to the Hawaiian bobtail squid symbiont *Vibrio fischeri* and the medically relevant pathogen *P. aeruginosa* PA14. This technique is generally applicable to biofilm-forming bacterial species and has significant potential to generate biological hypotheses regarding inter-related metabolic pathways influencing biofilm formation and dispersal.

## Results

### Spatial detection of c-di-GMP in *V. cholerae* biofilm colonies

Initially, we sought to test both positive mode and negative mode matrices for their ability to ionize c-di-GMP. For positive mode imaging, a 1:1 mixture of α-cyano-4-hydroxycinnamic acid (CHCA) and 2,5-dihydroxybenzoic acid (DHB) matrices were applied using a sieve method for bacterial colonies grown on agar media.^21^ We tested two other MALDI matrices that are known to facilitate ionization of nucleotides and nucleosides, 2′,4′,6′-trihydroxyacetophenone (THAP) and 3-hydroxypicolinic acid (HPA),^22,23^ in addition to CHCA and DHB in negative ionization mode and determined that the 1:1 CHCA:DHB matrix provided the best limit of detection for c-di-GMP detection (Methods, Fig. S1). As a proof of concept, we used two model *V. cholerae* El Tor strains to compare c-di-GMP production, one of which is well-established as an overproducer of c-di-GMP. The *V. cholerae* O1 El Tor A1552 wildtype strain produces a smooth phenotype when grown on LB agar, while the *V. cholerae* O1 El Tor A1552 rugose variant produces a wrinkled phenotype on LB agar due to an excess of biofilm production (Table 1). The *V. cholerae* rugose variant used here has been well-established as a biofilm producer due to the activity of the DGC VpvC, which has a tryptophan to arginine amino acid residue change as the result of a single nucleotide polymorphism compared to the wildtype strain, rendering it hyperactive.^24,25^

**Table 1:** Bacterial strains genotype and sources.

| Species | Strain | Genotype | Ref. | Strain Name in Manuscript |
| --- | --- | --- | --- | --- |
| *V. cholerae* | Fy_Vc_1 | *Vibrio cholerae* O1 El Tor A1552, wild type, Rif<sup>r</sup> | 24 | wildtype |
| *V. cholerae* | Fy_Vc_2 | *Vibrio cholerae* O1 El Tor A1552, rugose variant, Rif<sup>r</sup> | 24 | rugose variant |
| *V. cholerae* | Fy_Vc_1745 | Fy_Vc_2 ΔvpvC | 25 | RΔvpvC |
| *V. cholerae* | FY_VC_11823 | FY_VC_1, pFY4950, Rif Gent | this study | wildtype biosensor |
| *V. cholerae* | FY_VC_17439 | FY_VC_2, pFY4950, Rif Gent | this study | rugose variant biosensor |
| *V. fischeri* | MJM2773 | MJM1100 / pEVS143-MifA | 38 | DGC overexpression |
| *V. fischeri* | MJM3091 | MJM1100 / pEVS143-VF_0087 | 38 | PDE overexpression |
| *V. fischeri* | MJM4094 | MJM1100 / pRYI039 | 38 | wildtype |
| *P. aeruginosa* | PA14 pMMB | PA14 pMMB-gent | 17 | empty vector |
| *P. aeruginosa* | PA14 pMMB PA2200 | PA14 pMMB PA2200 | 17 | PDE overexpression |
| *P. aeruginosa* | PA14 pMMB PA1107 | PA14 pMMB PA1107 | 17 | DGC overexpression |
| *P. aeruginosa* | PA14 pMMB PA3702 | PA14 pMMB PA3702 | 17 | DGC overexpression |

Our initial results using wildtype *V. cholerae* and the rugose variant showed that c-di-GMP can be detected as both *m/z* 689.089 [M-H]^-^ and *m/z* 711.075 [M+Na-2H]^-^ (Fig. 1a). Both adduct ions of c-di-GMP show related distributions in the ion images of the wildtype and rugose variant *V. cholerae* strains, and the presence of these adducts was confirmed by the use of a c-di-GMP chemical standard applied by the dried droplet method to a piece of LB agar, which was prepared for MSI using the sieve method with 1:1 CHCA:DHB (Fig. 1a). We further confirmed the identity of c-di-GMP using MS/MS fragmentation on dried droplet samples of a c-di-GMP chemical standard compared to an extract of the *V. cholerae* rugose variant biofilm colony (Fig. 1b). When comparing the spatial c-di-GMP distribution between the wildtype and rugose variant, the rugose variant showed higher concentrations of c-di-GMP along the edge of the colony, whereas the wildtype *V. cholerae* produced c-di-GMP more diffusely throughout the colony (Fig. 1a). The 200 μm raster width better aligned with the rugosity observed in the optical images of the *V. cholerae* rugose variant, while the wildtype did not have distinct spatial differences in c-di-GMP detection at either 200 μm or 500 μm raster widths (Fig. 1a). We proceeded with the 200 μm raster width for all subsequent experiments as it gave adequate spatial resolution and could be used to image an entire biofilm colony (Fig 1). We further analyzed a *V. cholerae* rugose variant lacking the dominant DGC, VpvC, using MSI (RΔ*vpvC*, Fig. 1c). The RΔ*vpvC* mutant had a smooth colony morphology, which was reflected in the change in both intensity and distribution of c-di-GMP detected via MSI (Fig. 1c).

**Figure 1.**
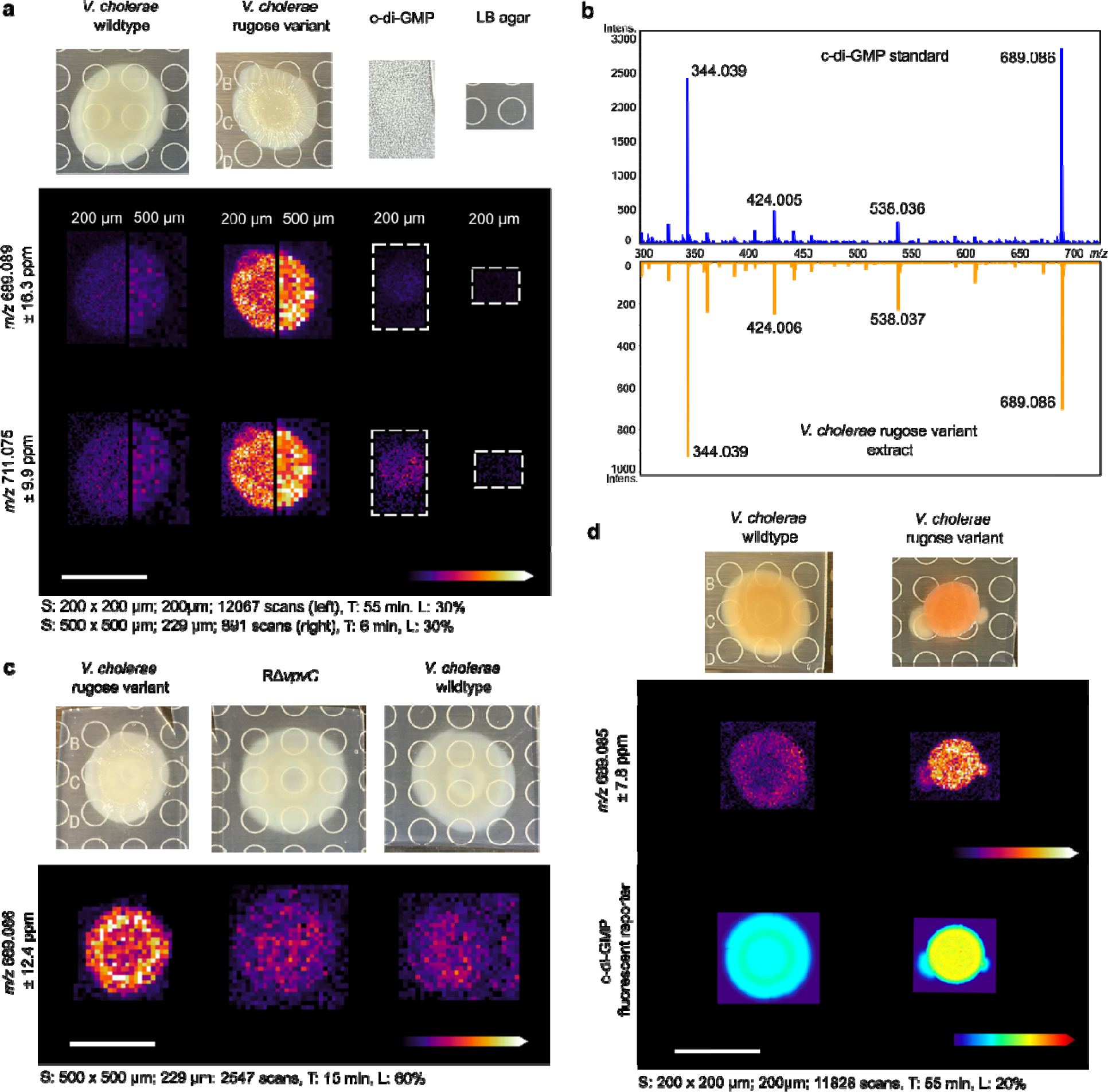
Spatial detection and validation of c-di-GMP in *V cholerae* wildtype and rugose variant strains. **a)** Ion images of the [M-H]^-^ (m/z 689.089) and [M-2H+Na]^-^(m/z 711.075) adducts of c-di-GMP in *V. cholerae* wildtype and rugose variant colonies. Data were acquired at both 200 μm and 500 μm raster to compare image resolution and sensitivity. An authentic standard of c-diGMP (5 μL, 100 n) was spotted on a sample of LB agar for comparison. **b)** MALDI-MS/MS spectrum of c-di-GMP compared to an extract of a *V. cholerae* rugose variant colony grown on LB agar. **c)** Ion images of c di-GMP in *V. cholerae* wildtype, rugose variant, and a rugose variant Δ*vpvC* (RΔvpvC). **d)** Ion images of c-di-GMP in *V. cholerae* wildtype and the rugose variant compared to the c-di-GMP specific reporter. Abundance of c-di-GMP is represented by heat maps showing the relative TurboRFP fluorescent signal in the same bacterial colonies. Spot raster; size; scan number (S), acquisition time (T), and laser po er (L) shown for each MSI experiment. All scales bars represent 1 cm.

### Orthogonal validation of c-di-GMP detection using a riboswitch biosensor

To validate the spatial distributions we observed with MSI, we used an orthogonal fluorescent riboswitch biosensor for c-di-GMP detection.^9,26^ We grew strains on thin agar to directly compare growth conditions across both methodologies, imaged the colonies using a fluorescent microscope, then excised the same bacterial colonies for MALDI-MSI analysis. To measure c-di-GMP in live cells prior to MSI, we used a dual fluorescent c-di-GMP specific reporter that we have previously validated to accurately measure changes in c-di-GMP.^26^ The fluorescent protein AmCyan encoded in this biosensor is produced constitutively and is used to normalize expression of TurboRFP, which is regulated by two c-di-GMP riboswitches to report intracellular c-di-GMP levels. In this reporter, we introduced an AAV degron to TurboRFP to improve spatial measurement of c-di-GMP levels. Here we represent c-di-GMP abundance within spot biofilms as a heat map of TurboRFP fluorescent intensity throughout the biofilm; AmCyan fluorescence was comparable between strains.

The corresponding images shown in Figure 1d indicate that both the MSI and fluorescent biosensor detection methods can accurately detect differences between the low and high c-di-GMP producing strains. However, the MSI images afforded a higher resolution in the *V. cholerae* rugose variant colonies and increased sensitivity in the wildtype *V. cholerae* colonies (Fig. 1).

### Production of c-di-GMP in rugose and smooth *V. cholerae* colonies varies over time

We sought to measure c-di-GMP levels over time in *V. cholerae* strains to observe how spatial changes in c-di-GMP production develop in wildtype *V. cholerae* and the rugose variant. All biofilm colonies were inoculated at the same time and three colonies each from the wildtype and rugose variant of *V. cholerae* were analyzed by MSI every 24 hours for four days. At each time point, the rugose variant colonies produced more c-di-GMP than the wildtype colonies (Fig. 2). This difference was most apparent at 24 hours, where c-di-GMP was barely detectable in the wildtype *V. cholerae* biofilm colonies. The c-di-GMP levels in the wildtype *V. cholerae* biofilms increased around 72 hours, but maintained a different spatial distribution around the edge of the colony, as opposed to the rugose variant which had the highest c-di-GMP levels in the center of the colony. (Fig. 2) We applied a Wilcox Rank Sum Test using the SCiLS MSI analysis software (Bruker Daltonics) as a statistical analysis to compare the ion intensity between the wildtype and rugose variants at each time point and found that these strains had significant differences in c-di-GMP detection at all four time points (Methods, Fig. 2).

**Figure 2.**
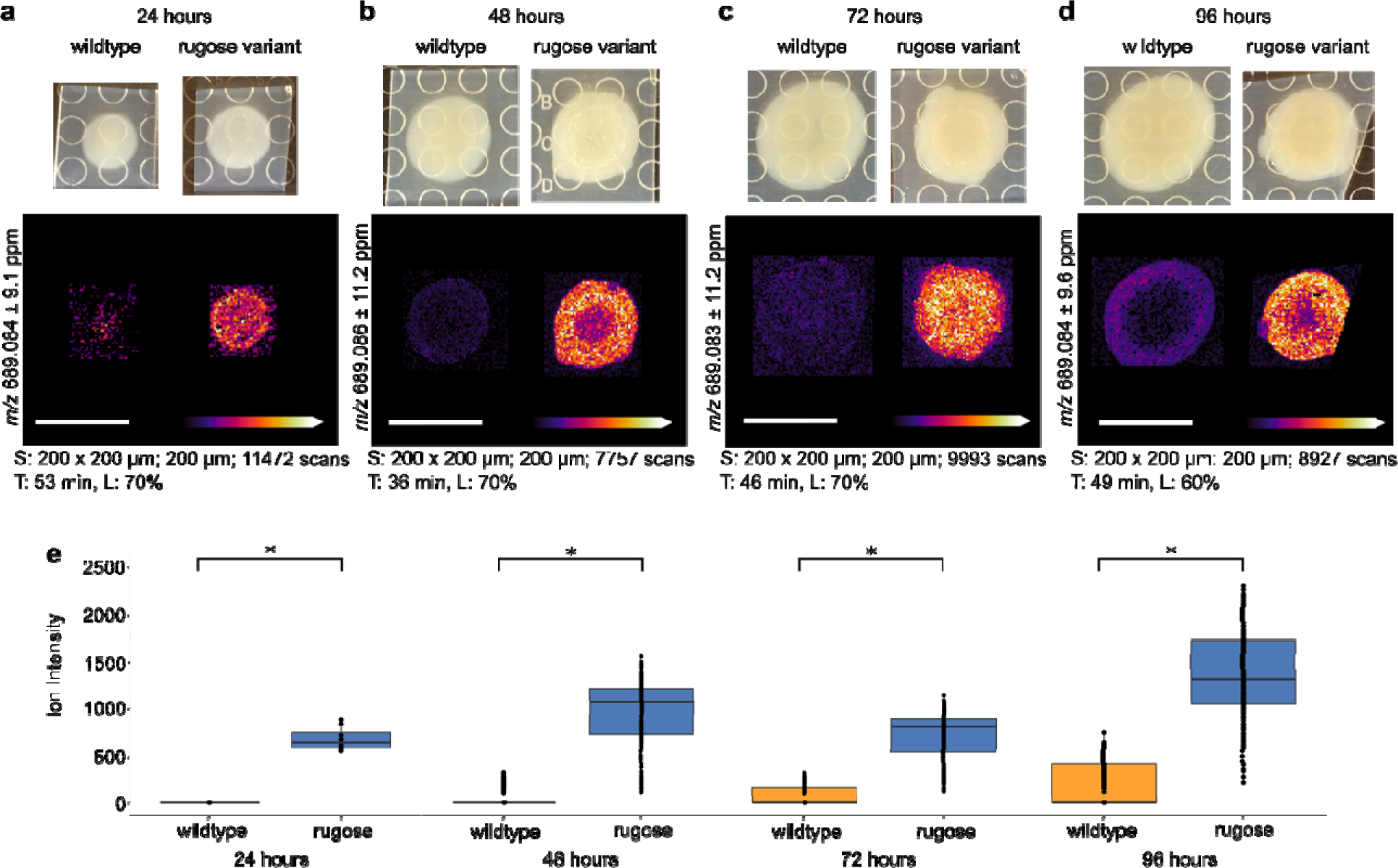
Comparison of c-di-GMP spatial distribution in *V. cholerae* colonies over time. Ion images and boxplots comparing ion intensity for *V. cholerae* wildtype and rugose variant strains after **a)** 24 hours, **b)** 48 hours, **c)** 72 hours, and **d)** 96 hours of growth. Spot raster; size; scan number (S), acquisition time (T), and laser power (L) shown for each MSI experiment. All scale bars represent 1 cm. **e)** Box plots comparing ion intensity between *V. cholerae* wildtype and rugose variants grown on each day. The box plots are shown on a single graph, however the data from each time point were acquired as separate experiments. The line within each box plot represents t e median, the edges of the box plots represent the upper and lower ends of the interquartile range and the whiskers represent the minimum and maximum quartiles. Outliers are shown beyond the whiskers of the box plots. Asterisks indicate statistical significance (p < 0.001) in a Wilcox Rank Sum test which was calculated using the SciLs MSI analysis software.

### Imaging c-di-GMP in a marine bobtail squid symbiont *Vibrio fischeri*

*V. fischeri*, another biofilm-producing relative in the *Vibrionaceae*, is the sole light organ symbiont of *Euprymna scolopes* (Hawaiian bobtail squid). The *V. fischeri* and *E. scolopes* relationship is a well-established model symbiosis that requires biofilm production by *V. fischeri* and for which c-di-GMP influences colonization behaviors ^6,27–30^. The symbiotic biofilm produced by *V. fischeri* is regulated differently from *V. cholerae* biofilm production ^28,31,32^, while cellulose production by *V. fischeri* is regulated by the c-di-GMP-responsive transcriptional regulator VpsR, a homolog of the *V. cholerae* master regulator of biofilm ^29,33–35^. This difference in biofilm regulation highlights the complexity and importance of studying the relative biological influence of c-di-GMP-mediated biofilm production in different contexts. To further validate MSI as an effective detection method for c-di-GMP, we applied well-established genetic manipulations to enzymes in the *V. fischeri* c-di-GMP pathway (Table 1).^36,37^ These two genetic alterations in the c-di-GMP pathway result in opposite c-di-GMP phenotypes when grown on agar,^38^ so we focused on the following *V. fischeri* ES114 derivatives to perform these studies: (i) a strain overexpressing *V. fischeri* PDE VF_0087^39^ from an IPTG-inducible vector and has smooth colony morphology (“PDE overexpression”; i.e., ES114/pEVS143-VF_0087), (ii) wildtype ES114 strain with the empty vector (“WT”; i.e., ES114/pRYI039), and (iii) a strain overexpressing *V. fischeri* DGC MifA^36^ that exhibits a wrinkled morphology (“DGC overexpression”; i.e., ES114/pEVS143-MifA) (Table 1, Fig. 3). We have shown previously that an analogous gradient of strains can be used to detect other compounds that are upregulated in biofilm colonies, and here we show that these methods can be expanded to other compounds and genetic contexts.^40^

**Figure 3.**
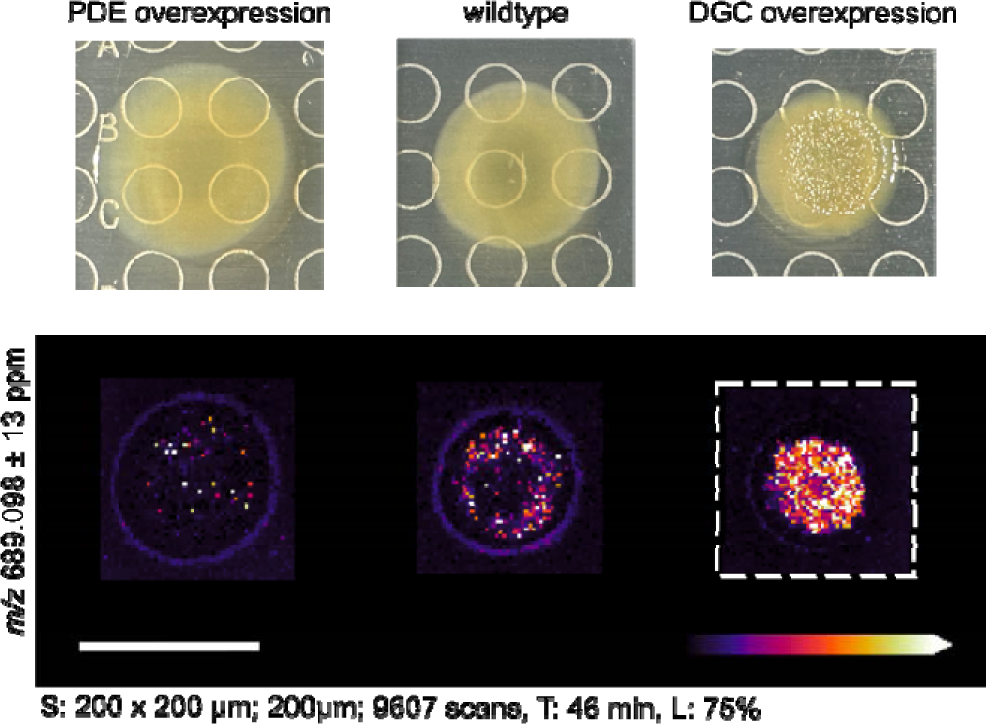
Comparison of c-di-GMP spatial distribution in *V. fischeri* colonies. The low c-di-GMP (PDE overexpression) and high c-di-GMP (DGC overexpression) strains contain a plasmid with an inducible promoter for the overexpression of the PDE VF_0087 and DGC MifA, respectively. The wildtype strain contains the vector control only. Spot raster; size; scan number (S), acquisition time (T), and laser power (L) shown for each MSI experiment. All scale bars represent 1 cm.

Using MSI we determined that the spatial distribution of c-di-GMP in *V. fischeri* colonies correlated to the wrinkles within these colonies, and the highest intensity signal across colonies assessed was associated with the DGC overexpression strain (Fig. 3, Fig. S5). Validating the strains used, the lowest intensity signal corresponded to the PDE overexpression strain, while a small amount of c-di-GMP was present in the wildtype (Fig. 3). Similarly to *V. cholerae*, higher c-di-GMP levels were detected throughout the DGC overexpression colony, whereas lower levels of c-di-GMP were detected near the edges of the wildtype and PDE overexpression strains (Fig. 3).

### Integration of untargeted metabolomics with c-di-GMP imaging in *P. aeruginosa* PA14

*P. aeruginosa* is an important opportunistic pathogen and prolific producer of specialized metabolites, especially those related to virulence. Biofilm production, virulence, and specialized metabolism are intricately connected in *P. aeruginosa* through various signaling pathways.^41,42^ We have previously used MSI to study changes in metabolite production by *P. aeruginosa* PA14 due to the presence of an exogenous conjugated bile acid, taurolithocholic acid.^43^ Here we used MSI to analyze *P. aeruginosa* PA14 biofilms for both c-di-GMP production in negative mode ionization, and some common classes of important *Pseudomonas* metabolites in positive mode ionization (Fig. 4). We applied this method to three *P. aeruginosa* PA14 strains, each containing an overexpression vector for a PDE (pMB-PA2200) or DGC (pMMB-PA1107, pMMB-PA3702) and an empty vector control (pMMB) (Table 1, Fig. 4, Fig. S6). The negative ionization results show a strong colony-associated signal for c-di-GMP in the DGC overexpression strains (PA1107, PA3702) compared to both the empty vector and PDE overexpression strain (PA2200) which show only background levels of c-di-GMP (Fig 4a, Fig. S6a,c). The strains used here were previously found to have differing virulence, biofilm phenotypes, and global levels of c-di-GMP due to the overexpression of specific DGCs or PDEs. However, the specific enzymatic contributions to the local pools of c-di-GMP could not be determined.^17^ Our results further validate the utility of our method to study local c-di-GMP production and spatial distribution in biofilm forming organisms under different genetic and environmental influences.

**Figure 4.**
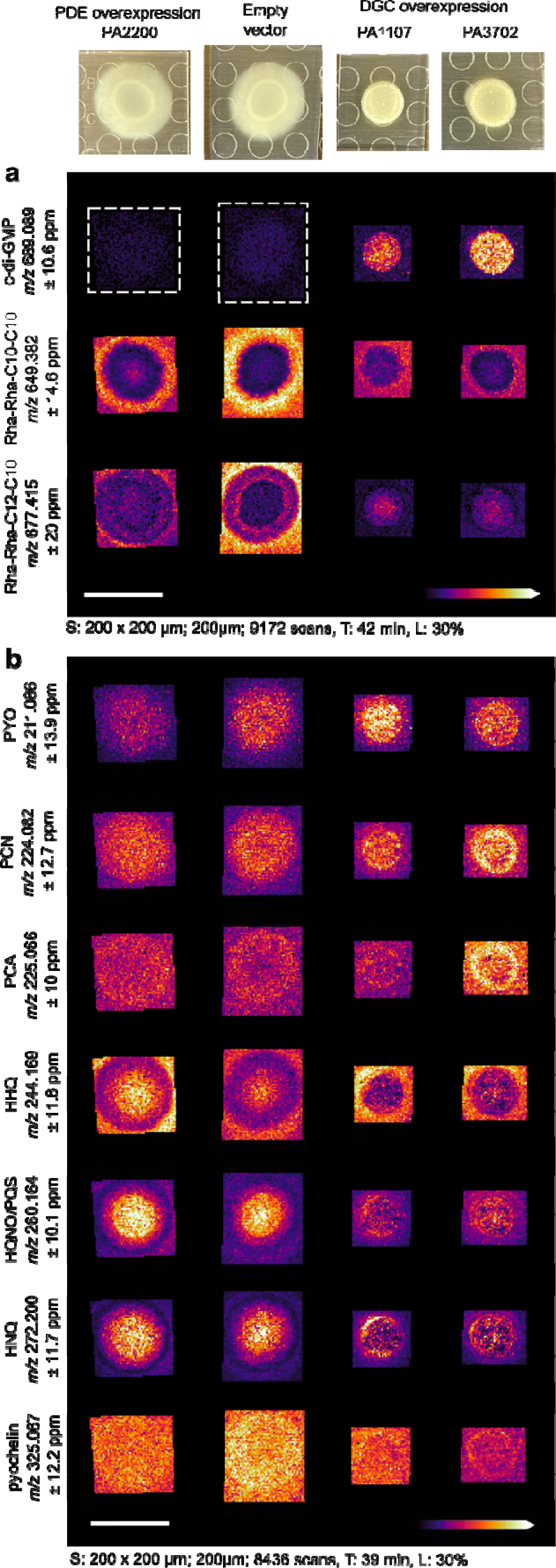
Ion images of c-di-GMP and other putatively identified metabolites in *P. aeruginosa* PA14. Ion images of metabolites detected in **a)** negative mode ionization and **b)** positive mode ionization. The following compound abbreviations are used: pyocyanin (PYO), phenazine-1-carboxamide (PCN), phenazine-1-carboxylic acid (PCA), *Pseudomonas* quinolone signal (PQS), 4-hydroxy-2-heptyquinoline-N-oxide (HQNO), 2-heptyl-4-quinolone (HHQ), and 4-hydroxy-2-nonylquinoline (HNQ) Table 2 shows the ppm error for all putatively identified compounds. Spot raster; size; scan number (S), acquisition time (T), and laser power (L shown for each MSI experiment. All scale bars represent 1 cm.

**Table 2:**
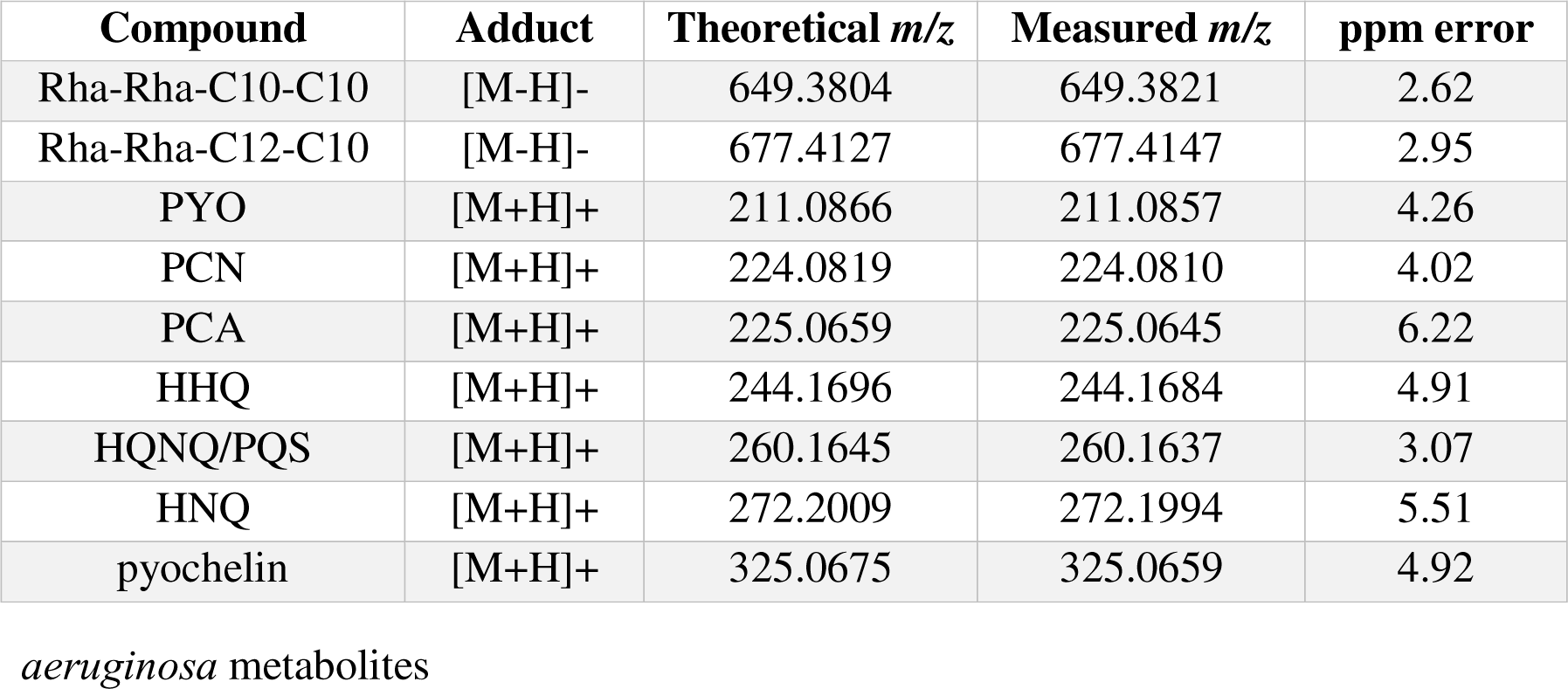
Mass error for detect ed *P*.

| Compound | Adduct | Theoretical $m/z$ | Measured $m/z$ | ppm error |
| --- | --- | --- | --- | --- |
| Rha-Rha-C10-C10 | [M-H]- | 649.3804 | 649.3821 | 2.62 |
| Rha-Rha-C12-C10 | [M-H]- | 677.4127 | 677.4147 | 2.95 |
| PYO | [M+H]+ | 211.0866 | 211.0857 | 4.26 |
| PCN | [M+H]+ | 224.0819 | 224.0810 | 4.02 |
| PCA | [M+H]+ | 225.0659 | 225.0645 | 6.22 |
| HHQ | [M+H]+ | 244.1696 | 244.1684 | 4.91 |
| HQNQ/PQS | [M+H]+ | 260.1645 | 260.1637 | 3.07 |
| HNQ | [M+H]+ | 272.2009 | 272.1994 | 5.51 |
| pyochelin | [M+H]+ | 325.0675 | 325.0659 | 4.92 |

*aeruginosa* metabolites

We then putatively identified other common specialized metabolites at the MS1 level in both negative and positive ionization modes from the same colony. Rhamnolipids are important biosurfactant molecules that are involved in mediating biofilm structure and dispersal.^44,45^ We detected two signals in negative mode that correspond to the rhamnolipids Rha-Rha-C10-C10 and Rha-Rha-C12-C10 (Fig. 4a). Both rhamnolipids were detected at a higher intensity in the empty vector strain compared to all PDE and DGC overexpression strains (Fig. 4a). This was expected as rhamnolipids are known to mediate cellular motility and are known to be anti-correlated with the production of phenazines such as pyocyanin (PYO) which are upregulated in biofilms.^46^ While we are not able to distinguish the isomeric rhamnolipid species using this method, we can identify distinct differences in the spatial distribution of different rhamnolipids in both low and high c-di-GMP conditions. Both the PDE overexpression strain, and the DGC overexpression strains had a decreased signal intensity for both rhamnolipids compared to the empty vector (Fig. 4a), however we can also see that the Rha-Rha-C12-C10 rhamnolipid was detected more within the biofilm colonies in all conditions compared to the Rha-Rha-C10-C10 rhamnolipid which appears to be primarily secreted from the biofilm colony in all conditions (Fig. 4a).

Phenazines are redox active molecules that are well-established as a microbial biofilm response to oxidative stress,^47^ and implicated in virulence in clinical isolates.^42^ We have previously shown how MSI can be used to interrogate changes in *P. aeruginosa* specialized metabolism due to both genetic mutations in the phz gene cluster and the presence of external chemical stimuli.^43^ Here our results show that phenazine production is slightly higher in the DGC overexpression strains compared to the empty vector control and the PDE overexpression strain (Fig. 4b), but overall the phenazine production varies slightly between each strain. While the changes detected here are minor, the consistent detection of these important molecules may serve as a benchmark for understanding global changes to *P. aeruginosa* metabolism in different genetic and environmental contexts.

We also detected the quinolone family of quorum sensing molecules using positive mode ionization (Fig. 4b). *Pseudomonas* quinolone signal (PQS), 4-hydroxy-2-heptyquinoline-N-oxide (HQNO), 2-heptyl-4-quinolone (HHQ), and 4-hydroxy-2-nonylquinoline (HNQ) are well established signaling molecules that we have previously identified in *P. aeruginosa* PA14.^43^ The isomeric species PQS and HQNO cannot be distinguished by our current MSI method (Fig. 4b), although we have previously used orthogonal methods to demonstrate that PQS and HQNO are produced in different spatial distributions in PA14.^43^ The detected quinolones exhibited a distinct spatial distribution where HHQ had higher intensity around the edge of the biofilm colony compared to HQNQ/PQS and HNQ which showed the highest ion intensity only within the biofilm colony in all conditions (Fig. 4b, Fig. S6b,d).

Pyochelin is a siderophore which is also implicated in virulence of *P. aeruginosa*. We have previously shown how pyochelin production is mediated by exposure to exogenous compounds such as bile acids.^43^ We found that the pyochelin signal was highest in the empty vector and the PDE overexpression strain, and lowest in the DGC overexpression strain pMMB-PA3702 (Fig 4b, Fig. S6b,d).

## Discussion

The signaling molecule c-di-GMP plays an important role in the complex coordination of bacterial metabolism throughout biofilm formation, maintenance, and dispersal. The accurate and sensitive detection of c-di-GMP in different contexts will inform how local and temporal changes in c-di-GMP concentrations control biofilm physiology. While many fluorescence-based techniques are in active development and expansion, we present a label-free MSI method for detecting c-di-GMP spatially in different organisms and environmental contexts. Here we show how this method can be applied to genetic mutants and natural isolates from both pathogenic and symbiotic contexts to compare spatial changes in local pools of c-di-GMP.

All of the model microorganisms presented here have genetic modifications specifically related to their c-di-GMP production. Biological turnover of c-di-GMP is complex and is likely controlled by local environmental sensing to improve adaptability and flexibility throughout the infection process. These local fluctuations in c-di-GMP production that influence biofilm phenotypes are relevant in both medical and environmental contexts. There is a need to interrogate the relative influence of c-di-GMP in microenvironments both via genetic and environmental manipulations, but also in host tissue contexts. Using the *E. scolopes* and *V. fischeri* model system, we have previously shown how MSI can be optimized on agar grown biofilms for further application to host tissues.^40^

*P. aeruginosa* is well-studied for its arsenal of small metabolite virulence factors, and we have highlighted some of the commonly studied compounds that are known to influence *P. aeruginosa* virulence and biofilm phenotypes.^41,43,48^ Our MALDI-MSI method allows for the simultaneous detection of c-di-GMP as well as other specialized metabolites in both positive and negative ionization mode from a single sample by repeated laser irradiation in two experiments (Fig. 4). While the global levels of c-di-GMP have been well-studied in relation to genetic and environmental manipulation, these measurements do not always correlate with the phenotypic alterations observed in different DGC or PDE mutants.^17^ The local contribution of different DGC and PDE enzymes is likely to be a major influence in the altered biofilm phenotypes that are observed. MALDI-MSI detection of c-di-GMP in bacterial colonies allows for the efficient observation of local changes to specialized metabolite pathways in correlation with changes in c-di-GMP levels. The versatility of this method allows for the spatial correlation of c-di-GMP with various specialized metabolites in genetically engineered strains as well as clinical isolates and has the potential to be optimized for the direct application to host tissues.

### Online Methods

#### Bacterial strains and growth conditions

All bacterial strains used in this study are described in Supplementary Table 1. All *V. cholerae* strains were grown in Bacto LB media (10g NaCl, 10 g tryptone, 5 g yeast extract, pH 7.5 in 1 L DI H2O). The *V. cholerae* (biosensor) strains and *P. aeruginosa* PA14 strains were grown in LB media containing 15 µg/mL gentamicin. *V. fischeri* was grown in LBS (LuriaBertani salt: 20 g NaCl, 10 g tryptone, 5 g yeast, 50 mL 1 M Tris, pH 7.0 in 1 L DI H_2_O) agar with 100 µg/mL kanamycin.

All bacteria were plated on 1.5% agar media and grown overnight at 30 °C. A colony from the plate was then used to inoculate a 5 mL liquid culture of and grown overnight at 30 °C while shaking at 225 rpm. Overnight liquid cultures were normalized to an OD_600_ of 0.1 by diluting in sterile media and 5 μL of inoculum was spotted on thin agar plates (3 mL of agar in a 60 mm plate) and incubated for up to 96 h at 30 °C (*V. fischeri* was grown at room temperature). For *P. aeruginosa* PA14 strains, 100 µM IPTG was added to the thin agar plates for plasmid induction in the bacterial colonies that were analyzed by MALDI-MSI.

#### Colony sample preparation for MSI

Following 96 h of growth, colonies were excised from the agar plates using a razor blade and transferred to an MSP 96-target ground-steel target plate (Bruker Daltonics). Two additional pieces of sterile agar were excised and transferred to the target plate as controls. An optical image of the colonies on the target plate was taken prior to matrix application. A 53 μm stainless steel sieve (Hogentogler Inc.) was used to coat the steel target plate and colonies with MALDI matrix. The MALDI matrix used for the analysis was a 1:1 mixture of recrystallized δ-cyano-4-hydroxycinnamic acid (CHCA) and 2,5-dihydroxybenzoic acid (DHB) (Sigma). The plate was then placed in an oven at 40 °C for approximately 4 h or until the agar was fully desiccated. After 4 h, excess matrix was removed from the target plate and sample with a stream of air. A chemical standard of c-di-GMP (1uL, 100nM) was spotted using a dried droplet method, onto one of the desiccated agar control samples. Another optical image was taken of the desiccated colonies on the target plate.

#### MALDI-MSI analysis of bacterial colonies

To determine if our standard MALDI matrix composition would be sufficient to ionize c-di-GMP in positive or negative ionization mode, we compared 1:1 CHCA:DHB matrix to two additional matrices that have been used for MALDI analysis of oligonucleotides using a dried droplet method with a c-di-GMP chemical standard. We tested a 1:1 CHCA:DHB mixture, CHCA alone, 2′,4′,6′-Trihydroxyacetophenone (THAP), and 3-hydroxypicolinic acid (HPA) in positive and negative mode. The positive ionization mode did not provide adequate c-di-GMP ionization regardless of the MALDI matrix used. This is likely due to the fact that typical acidic additives such as TFA cannot be added to matrices added using our dry sieve matrix application method.^1^ In negative mode, we found that c-di-GMP ionizes well when CHCA is used alone or in combination with DHB, while the THAP and HPA matrices did not induce sufficient ionization at low c-di-GMP concentrations (Fig. S1). We opted to move forward using 1:1 CHCA:DHB for MSI analysis in negative mode, as it provided sufficient ionization for c-di-GMP, and was already optimized for the sieving method of matrix application and sample drying for MSI. These sample conditions may also be used on bacterial samples that are dried and prepared using a spray method and further modifications can be made such as using CHCA alone, or using acidic additives to enhance positive mode ionization of c-di-GMP. We tested both 500 μm and 200 μm raster widths on the bacterial biofilm colonies initially to determine the proper resolution for comparing the spatial distribution of c-di-GMP (Fig. 1a). The 200 μm raster width provided a clearer image that aligned well with the biofilm structures observed in the optical images, so we moved forward using the 200 μm raster width in all subsequent experiments. While these raster widths work well for the agar-grown biofilm colonies analyzed in this study, this parameter can easily be adjusted for other preparations of biofilms (i.e. cryosectioning) as well as host tissue analysis to achieve higher resolution images of *in situ* biofilm infections.

The desiccated bacterial colonies and agar controls were then analyzed using a MALDI mass spectrometer (Bruker timsTOF fleX qTOF mass spectrometer). Imaging mass spectrometry data was acquired using timsControl v 4.1 and flexImaging software v. 7.2. The data were collected using the mass range 300-800 Da in negative mode and 100–1500 Da in positive mode. Data were acquired at 200 μm (200 × 200 μm laser size) or 500 μm (229 × 229 μm laser size) spatial resolution using the M5 defocus laser setting. Each raster point was acquired using 200 laser shots at 1,000 Hz in all experiments. Other details regarding the acquisition parameters are indicated in each figure using the ‘SMART’ standardized nomenclature.^49^ Step size, spot size, and scan number (S), molecular identification (M), annotations (A), mass resolution (R), time of acquisition (T) are provided. All molecular identifications (M) were done to the MS1 level or using a commercial standard, all annotations (A) were targeted and no database searching was used and the mass resolution (R) in all experiments was 65,000 FWHM at *m/z* 1222. Because these three parameters were consistent throughout all experiments, we only indicate step and spot size (S), time of acquisition (T) and additionally, the laser power (L) used in each experiment on each figure. Table 2 indicates the MS1 level ppm error of the putatively identified compounds in Figure 4 and Figure S6. The ppm range shown in each ion image represents the bin width used to generate the ion image. The mass spectrometer was calibrated using phosphorus red as a calibrant. All raw data are available in MassIVE (massive.ucsd.edu) under the doi: doi:10.25345/C5RV0DB1W

#### Bacterial colony extraction for MALDI-MS/MS

A standard dried droplet technique for spotting chemical samples for MALDI analysis involves mixing 1 µL of a dissolved sample with 1 µL of a saturated solution of MALDI matrix and spotting the mixture onto a MALDI target plate. This method was used for all chemical standards of c-di-GMP dissolved in water and mixed with a 1:1 mixture of CHA:DHB dissolved in 78:22 acetonitrile:H_2_O.

In order to use MS/MS fragmentation to identify c-di-GMP in the bacterial biofilm produced by the *V. cholerae* rugose variant, we took a bacterial colony grown on thin agar for four days and scraped the entire colony into a saturated mixture of 1:1 CHCA:DHB in 1:1 methanol:H_2_O. We then vortexed the sample, centrifuged (13,000 rpm, 2 min), and spotted this mixture directly onto the MALDI target plate for analysis. MALDI MS/MS data were acquired in negative ionization mode using a total of 1000 laser shots at 1000 Hz, selecting for *m/z* 689.0866 with a mass isolation window of 0.10 Da and a collision energy of 40 eV.

#### Data Analysis

All ion images were generated using SCiLS lab software version 2023c pro (Bruker Daltonics). The boxplot data were generated by exporting the sample regions from SCiLS for import into Metaboscape v. 2022b (Bruker Daltonics). Peaking picking and thresholding was performed in metaboscape and a csv file was exported containing all of the mass spectra data for the *m/z* feature corresponding to the [M-H]^-^ ion of c-di-GMP (*m/z* 689.09). Boxplots were generated in R using the integrated ion intensity values of the [M-H]^-^ ion of c-di-GMP across the scans (raster points) in each ion image. The Wilcox Rank Sum test was run in SCiLS which will generate an exact p-value when the result is > 0.001, however all p-values calculated in this study were determined to be < 0.001 using SCiLS.

#### Microscopy for biosensor imaging

Colonies were prepared as previously described for MALDI-MS/MS analysis using the strain harboring the c-di-GMP specific biosensor. These were grown on 1.5% agar LB plates containing gentamicin and grown for 72 hours. Colonies were imaged using a Zeiss AxioCam HRm with a 1x/.025 NA lens on an Axiozoom V.16 stand. Amcyan was measured with an excitation of 470/40, a 495 dichroic mirror, and an emission of 525/50. TurboRFP was measured with an excitation of 550/25, a 570 dichroic mirror of 570, and an emission of 605/70. Acquisition parameters were identical between samples for comparison of fluorescent intensity. Images were analyzed using Zen Blue software. Quantification of c-di-GMP was then performed in parallel using MALDI-MS/MS.

#### Safety Note

Some of the microbes used in this research are considered biosafety level 2 and are opportunistic pathogens. We prepare samples in a biosafety cabinet to minimize exposure and once coated with matrix and dried, we consider the pathogens safe to handle, as they are no longer viable.

## Acknowledgements

This work was supported by the National Institute of Allergy and Infectious Diseases of the NIH under Award Number R01AI102584 (FHY), the National Institute of General Medical Sciences of the NIH award R35GM148385 (MJM), and by National Science Foundation grants IOS-2220510 (LMS) and IOS-2220511 (MJM), UCSC Startup funds (LMS). We thank Benjamin Abrams, UCSC Life Sciences Microscopy Center RRID:SCR_021135, for technical support during fluorescent imaging and analysis. We thank Vincent Lee from the University of Maryland for supplying the PA14 strains and Jing Gao for assistance in preparing the ES114 strains.

