## Supplementary material for "A label-free approach for relative spatial quantitation of c-di-GMP in microbial biofilms": SI

^5^ Current Address: Department of Microbiology and Immunology, University of Minnesota Medical School, Minneapolis, MN 55455

| **Figure S1.** MALDI-MS spectra of a commercial c-di-GMP standard………………….. | S2 |
| --- | --- |
| **Figure S2.** Replicates for *V. cholerae* wildtype and rugose variants……………………. | S3 |
| **Figure S3.** Replicate data for MALDI-MSI and fluorescent microscopy of *V. cholerae.* | S4 |
| **Figure S4.** Comparison of c-di-GMP spatial distribution in *V. cholerae* over time……. | S5 |
| **Figure S5.** Replicate data for MALDI-MSI of *V. fischeri* …..…………………………. | S6 |
| **Figure S6.** Replicate data for MALDI-MSI of *P. aeruginosa* PA14…………………… | S7-8 |

**Figure S1**. MALDI-MS spectra of a commercial c-di-GMP standard crystallized with four different MALDI matrices, **a)** α-cyano-4-hydroxycinnamic acid (CHCA) alone, **b)** a 1:1 mixture of CHCA and dihydroxybenzoic acid (DHB), **c)** 2′,4′,6′-Trihydroxyacetophenone (THAP), and **d)** 3-hydroxypicolinic acid (HPA)

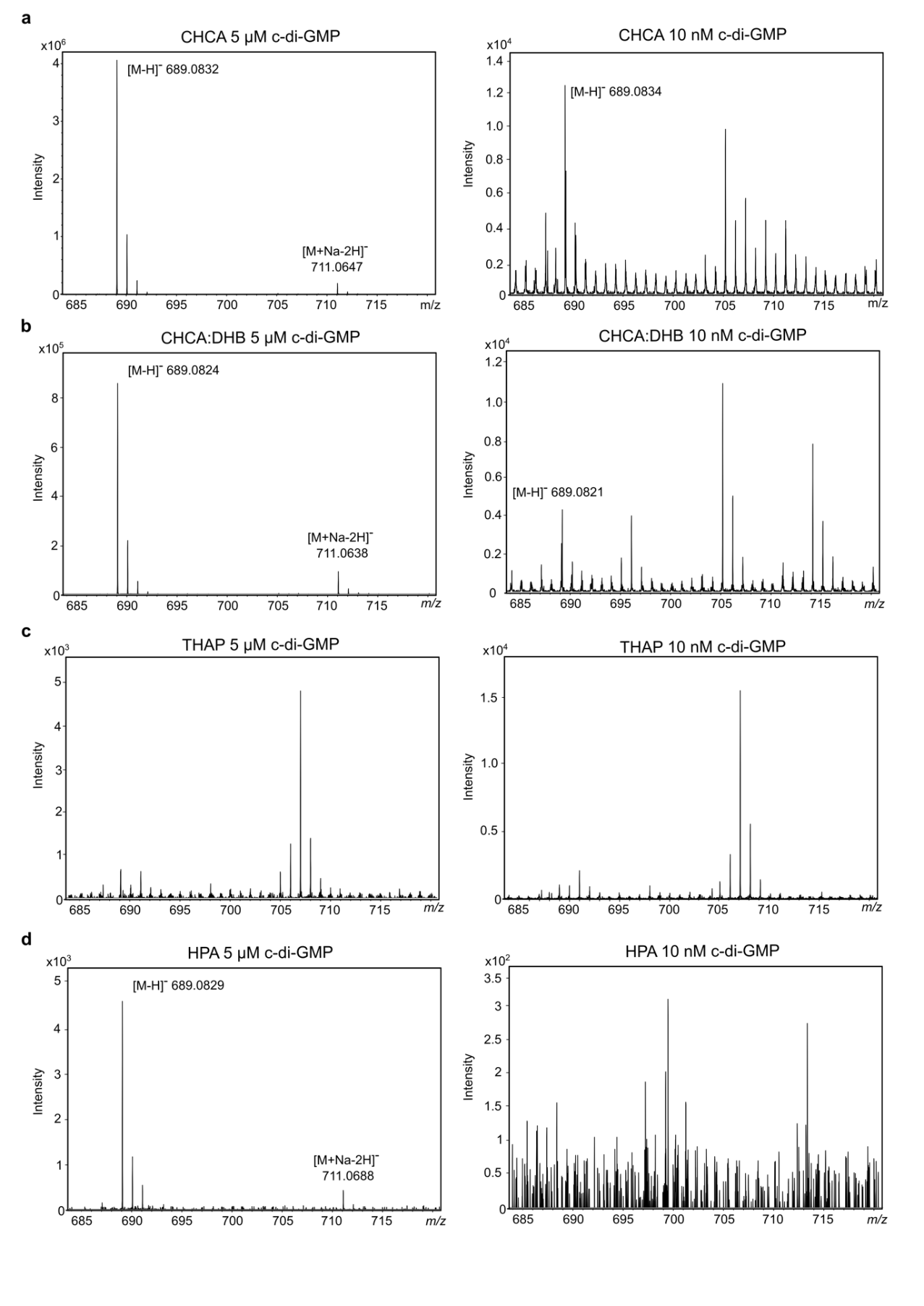

**Figure S2.** Replicate data for MALDI-MSI of *V. cholerae* in Figure 1. **a-b)** Ion images of replicates for *V. cholerae* wildtype and rugose variants from Fig. 1a. **c-d)** Ion images of replicates for *V. cholerae* wildtype, rugose variant, and R∆vpvC from Fig. 1c. Spot raster; size; scan number (S), acquisition time (T), and laser power (L) shown for each MSI experiment. All scale bars represent 1 cm.

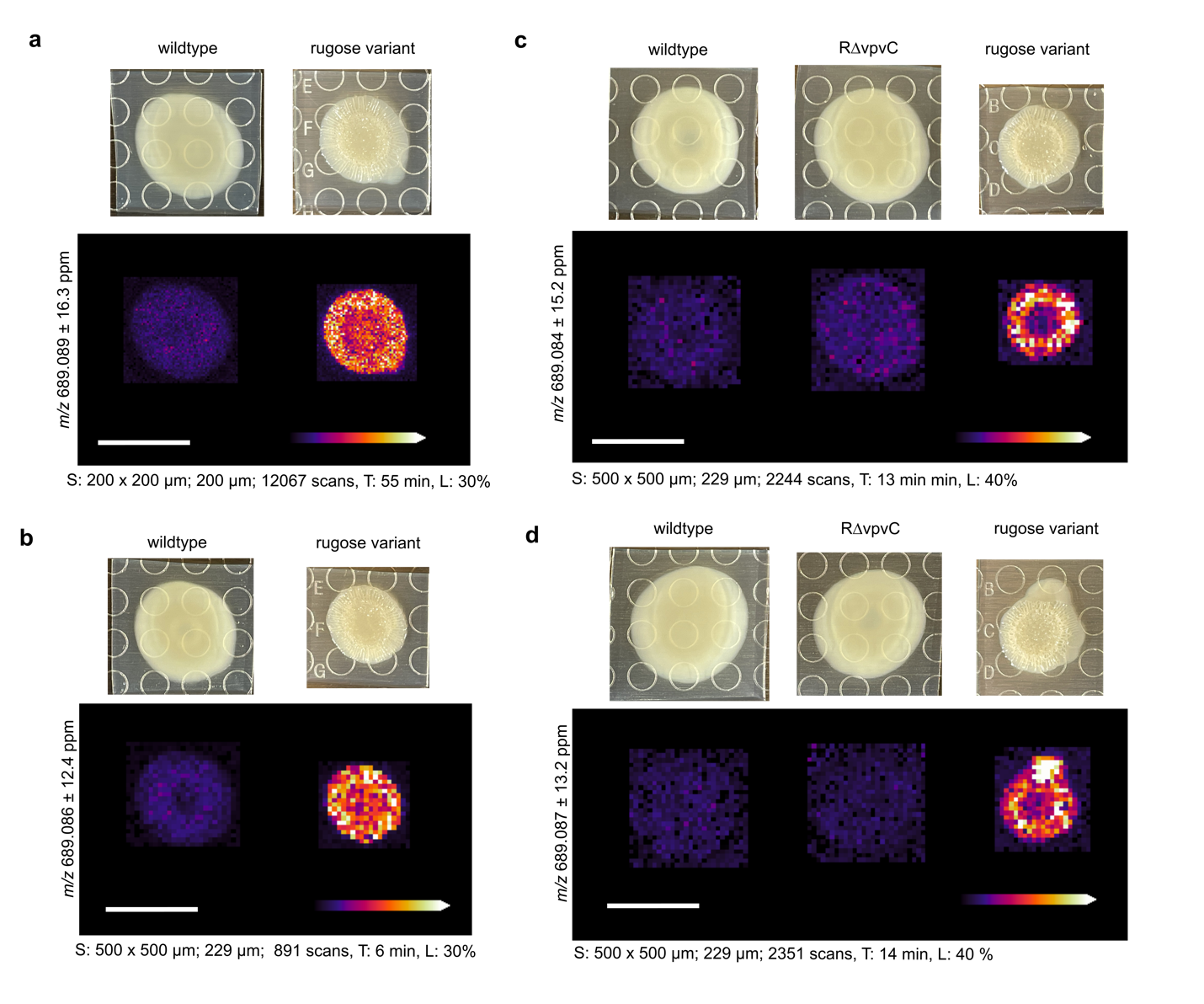

**Figure S3.** Replicate data for MALDI-MSI and fluorescent microscopy of *V. cholerae* in Figure 1d. Ion images of c-di-GMP in *V. cholerae* wildtype and the rugose variant compared to the c-di-GMP specific reporter. Abundance of c-di-GMP is represented by heat maps showing the relative TurboRFP fluorescent signal in the same bacterial colonies. Spot raster; size; scan number (S), acquisition time (T), and laser power (L) shown for each MSI experiment. All scale bars represent 1 cm.

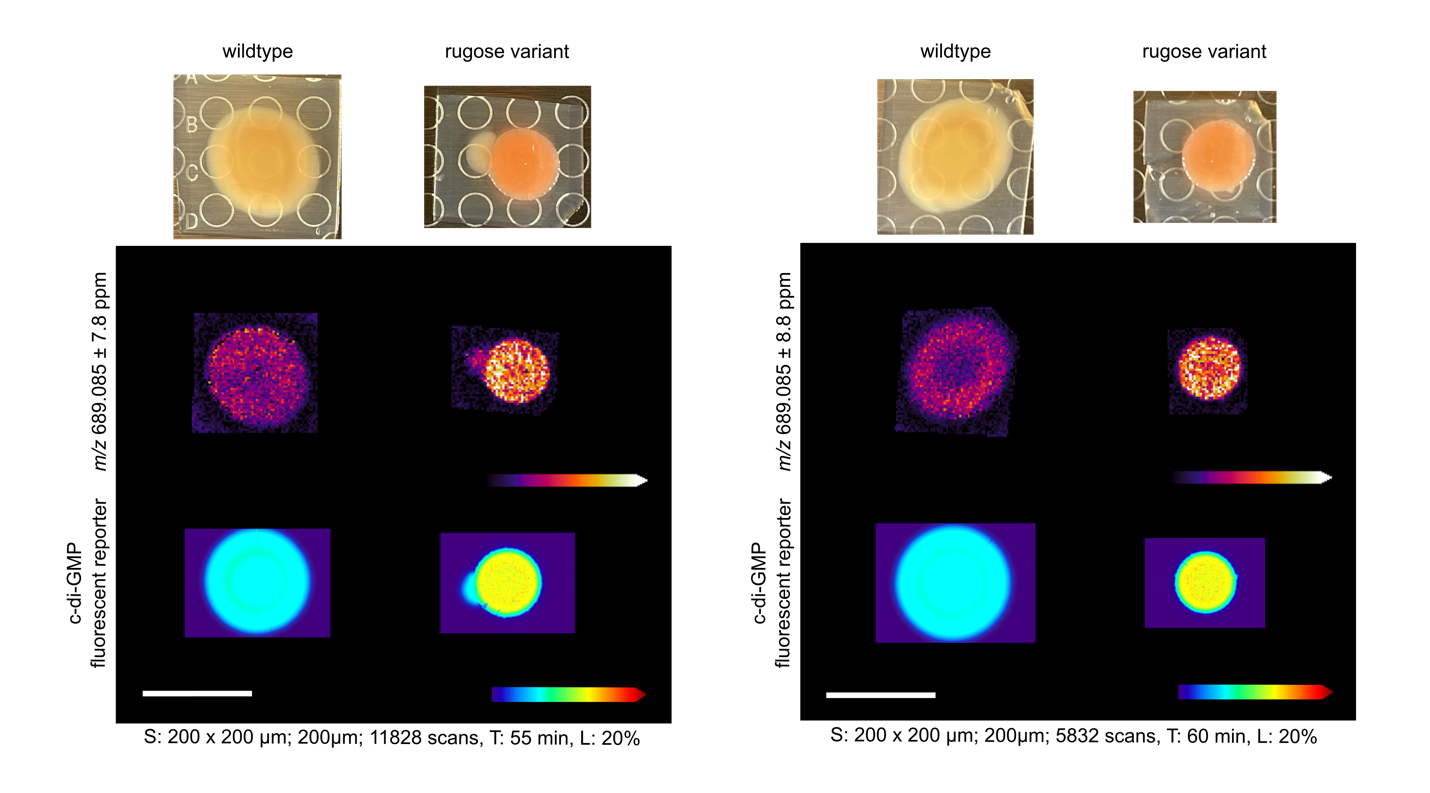

**Figure S4.** Replicate data for MALDI-MSI of *V. cholerae* in Figure 2. Comparison of c-di-GMP spatial distribution in *V. cholerae* colonies over time. Ion images for *V. cholerae* wildtype and rugose variant strains after **a)** 24 hours, **b)** 48 hours, **c)** 72 hours, and **d)** 96 hours of growth. Photos in d) show the biofilm colonies after MALDI matrix application and drying. Spot raster; size; scan number (S), acquisition time (T), and laser power (L) shown for each MSI experiment. All scale bars represent 1 cm.

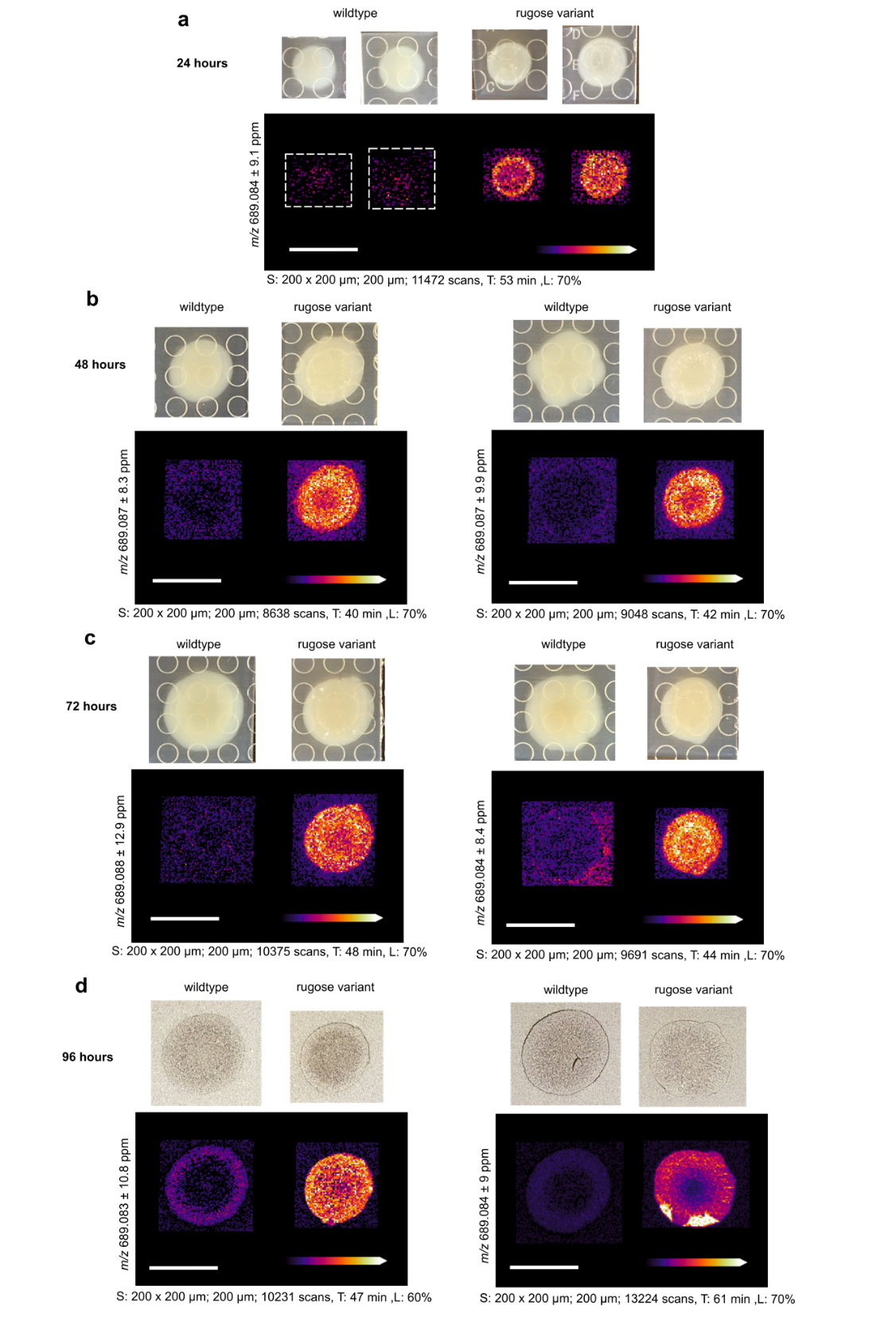

**Figure S5.** Replicate data for MALDI-MSI of *V. fischeri* in Figure 3. Ion images of c-di-GMP in three strains of *V. fischeri.* The low c-di-GMP (PDE overexpression) and high c-di-GMP (DGC overexpression) strains contain a plasmid with an inducible promoter for the overexpression of the PDE VF_0087 and DGC MifA, respectively. The wildtype strain contains the vector control only. Spot raster; size; scan number (S), acquisition time (T), and laser power (L) shown for each MSI experiment. All scale bars represent 1 cm.

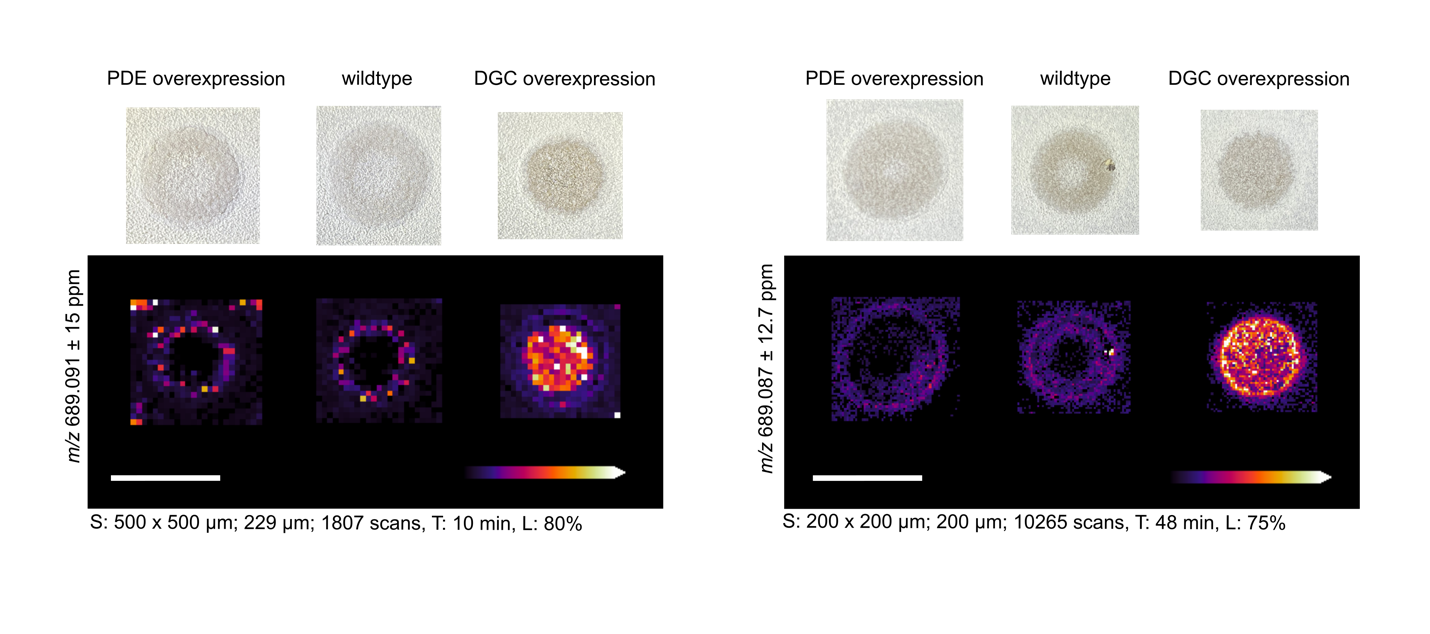

**Figure S6.** Replicate data for MALDI-MSI of *P. aeruginosa* PA14 in Figure 4 showing ion images of c-di-GMP and other putatively identified metabolites in *P. aeruginosa* PA14. Ion images of metabolites detected in **a)** negative mode ionization and **b)** positive mode ionization of one replicate and **c)** negative mode ionization and **d)** positive mode ionization of another replicate. The following compound abbreviations are used: pyocyanin (PYO), phenazine-1-carboxamide (PCN), phenazine-1-carboxylic acid (PCA), *Pseudomonas* quinolone signal (PQS), 4-hydroxy-2-heptyquinoline-N-oxide (HQNO), 2-heptyl-4-quinolone (HHQ), and 4-hydroxy-2-nonylquinoline (HNQ) Table 2 shows the ppm error for all putatively identified compounds. Spot raster; size; scan number (S), acquisition time (T), and laser power (L) shown for each MSI experiment. All scale bars represent 1 cm.

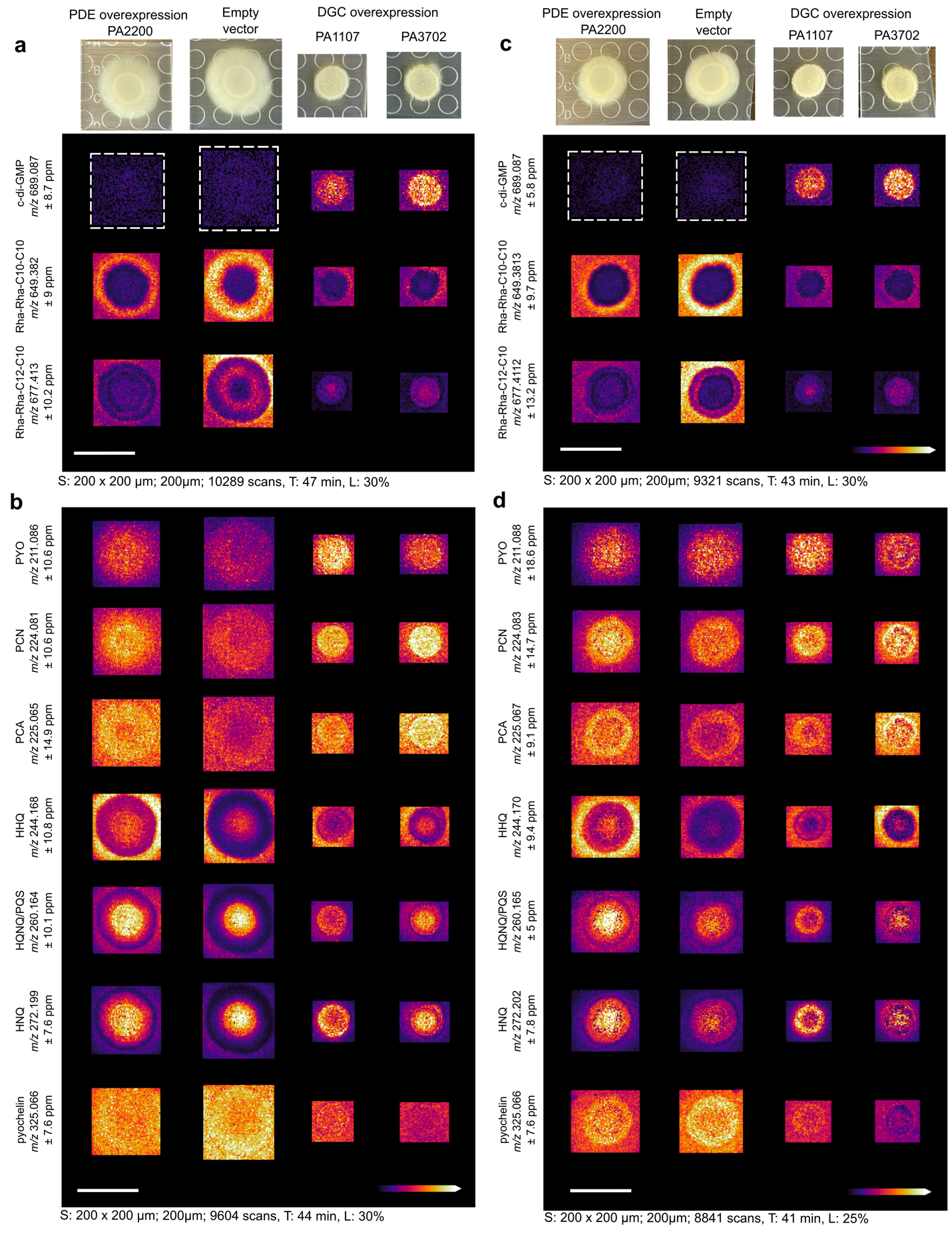
